# The SRG RAT® supports human cell xenotransplantation through enhanced tumor microenvironment interactions

**DOI:** 10.1101/2025.03.27.645250

**Authors:** Caitlin M. O’Connor, Kaitlin P. Zawacki, Jessica R. Durrant, Kelsey Barrie, R. Grace Walton, Dana King, Gabrielle Hodges Onishi, Fallon K. Noto, Matthew Hinderman, Diane Begemann, Taylor Branyan, Karah Panwell, Analisa DiFeo, Michael J. Schlosser, Goutham Narla

**Author notes:** To whom correspondence should be addressed: Goutham Narla, Department of Internal Medicine, Division of Genetic Medicine, University of Michigan, Ann Arbor, MI 48109.

## Abstract

The use of immunodeficient mice for human tumor engraftment is an essential model of human cancer, with uses ranging from basic science to translational research. However, low engraftment rates, slow growth, and smaller tumor volumes can be limitations. Previously, we reported a highly immunodeficient rat strain with the functional deletion of both the *Rag2* and *Il2rg* genes on the Sprague-Dawley background (**SRG RAT**^®^), which lacks B, T, and NK cells. Here, we subcutaneously engrafted two cell-derived xenograft (CDX) and seven patient-derived xenograft (PDX) models, including prostate, lung, ovarian, and uterine cancer models, into the SRG rat or NSG mouse models and tracked tumor growth. In all cases, the engraftment and tumor growth rates were better supported in the SRG rat compared to the NSG mouse. Interestingly, the SRG rat is not more immunocompromised than the NSG mouse, suggesting alternative mechanisms leading to the supportive growth in the SRG rat. Therefore, we explored potential differences in the tumor microenvironment (TME) between models grown in the two host animals. Lung PDX models grown in SRG rats showed enhanced formation of vasculature and stroma and were morphologically more consistent with the originating patient tumors. IHC analysis of the NCI-H660 CDX model showed differences in the tumors’ stroma, vasculature, and macrophages when grown in the two host species. Single-cell spatial imaging of engrafted tumors showed upregulation of the human *CXCL2* and *TCIM* in NCI-H660 tumors grown in the SRG rat versus the NSG mouse, both of which have been linked to poor prognosis in cancer. Combined, our data demonstrate that the SRG rat supports the growth of multiple human cancer types and displays enhanced tumor microenvironment interactions compared to NSG mice.

## Introduction

Our ability to model human cancer using animal models has improved dramatically over the past decades. Each model has advantages and disadvantages, making model selection a critical decision for every investigator. The use of immunodeficient mice for human tumor engraftment is an essential model of human cancer, with uses ranging from basic science to translational research (1–6). However, drug metabolism often differs between mice and humans, making them inferior models for pharmacokinetics, pharmacodynamics, and toxicology studies for most drug classes (7). Additionally, size-based advantages of rats include increased blood and tissue collection and easier surgical manipulations (8–11). As such, rats are the preferred rodent for preclinical studies due to their size and increased similarities to human drug metabolism (7). Use of mice is further limited by low tumor engraftment rates and slow tumor growth for many tumor models. Previously, we reported a new immunodeficient rat strain developed by Hera BioLabs, with the functional deletion of both the Rag2 and Il2rg genes on the Sprague-Dawley background (SRG RAT^®^), leading to loss of B, T, and NK cells (12). We showed that the SRG rat supports robust growth of multiple patient-derived xenograft (PDX) and cell-derived xenograft (CDX) models that have low take rates or poor growth in NSG mice (12). Thus, the SRG rat offers the potential for improved cancer xenografts in a species with advantages over mice.

The tumor microenvironment (TME) consists of tumor cells and neighboring stromal cells, which include endothelial cells, pericytes, fibroblasts, macrophages, and other immune cells (13–15). Stromal cells provide nutrition and structural support for tumors, and tumor-stroma interactions mediate tumor establishment, growth rate, and metastasis via cytokine signaling, growth factor signaling, and extracellular matrix interactions (16–18). SRG rats have been reported to be more permissive to engraftment of human cells compared to NSG mice, but the mechanisms underlying the supported growth are not well understood (8, 12, 19, 20). Notably, the SRG rat is not more immunocompromised than the NSG mouse, suggesting alternative mechanisms leading to enhanced tumor growth in the SRG rat. The current experiments were conducted to expand the number of xenograft models tested for direct comparison of tumor growth between the SRG and NSG animals and to further characterize TME in these tumors to better understand why xenotransplantation into the SRG rat enhances engraftment rates and tumor growth.

To this end, we compared tumor growth in SRG rats versus NSG mice using seven PDX models including lung, uterine, and ovarian cancers. We also compared tumor growth with EFO-27 ovarian carcinoma cells and NCI-H660 epithelial neuroendocrine prostate cancer cells. Next, we compared histopathology of lung adenocarcinoma PDXs grown in SRG rats or NSG mice to their original patient tumors. Lastly, we analyzed NCI-H660 tumor cell response to SRG versus NSG hosts using histopathology, immunohistochemistry (IHC), immunofluorescence (IF), and the 10X Xenium platform for spatially resolved single-cell RNA sequencing of the human tumor cells (21). Combined, our data demonstrate that the SRG rat supports the growth of multiple human cancer types and displays tumor morphology similar to patient tumors and enhanced TME interactions compared to NSG mice, highlighting the potential of the SRG rat to be a valuable model of human cancer.

## Results

### SRG rats support the growth of multiple PDX and CDX models over conventional immunodeficient mice

To understand why xenotransplantation into the SRG rat enhances engraftment rates and tumor growth compared to conventional immunodeficient mice, we engrafted seven patient-derived xenograft (PDX) models of lung (TM00233 and J000096652), ovarian (PDX111 and PDX133), or uterine (J0001123358, TM0001, and TM00274) origin and one cell derived xenograft (CDX) ovarian cancer model (EFO-27) in SRG rats or NSG mice and tracked tumor growth. PDX models were maintained in NSG mice, and when the tumors were large enough, they were harvested, sectioned, and serially implanted into recipient SRG rats or NSG mice. Tumor preparation and implantation were done at the same time and fragments of the same size were used for both host species (8 mm^3^). Tumors were measured twice weekly using a caliper to estimate the tumor volume.

Both lung adenocarcinoma PDX models displayed more robust tumor growth in the SRG rat (**Figure 1A and B**, **Table 1**). Interestingly, only the J000096652 model showed an increase in the engraftment rate in the SRG rat (**Figure 1B and Table 1**), and both species showed a similar time to engraftment (**Supplemental Figure 1**). The uterine endometrioid PDX model J000112358 showed earlier tumor growth in two of the NSG animals but established more robust growth in the SRG rat (**Figure 1C, Supplemental Figure 1**). The uterine sarcoma model TM0001 and uterine leiomyosarcoma model TM000274 both showed earlier engraftment times and more robust tumor growth in the SRG rats compared to NSG mice (**Figure 1D and E**, **Table 1, and Supplemental Figure 1**). TM0001 also showed a higher engraftment rate in SRG animals (**Table 1**). The ovarian cancer models PDX111, a high-grade serous ovarian carcinoma, and PDX133, an ovarian carcinosarcoma, both displayed more robust tumor growth in the SRG rats (**Figure 1F and G**). Additionally, these models had a 100% engraftment rate in the SRG rat (2/2), where PDX133 showed a 60% engraftment rate and PDX111 showed a 20% engraftment rate in the NSG mice (**Figures 1F and G**, **Table 1**). Finally, growth of the EFO-27 tumor fragments showed similar engraftment rate and growth between both host species (**Figure 1H**, **Table 1, and Supplemental Figure 1**).

**Figure 1:**
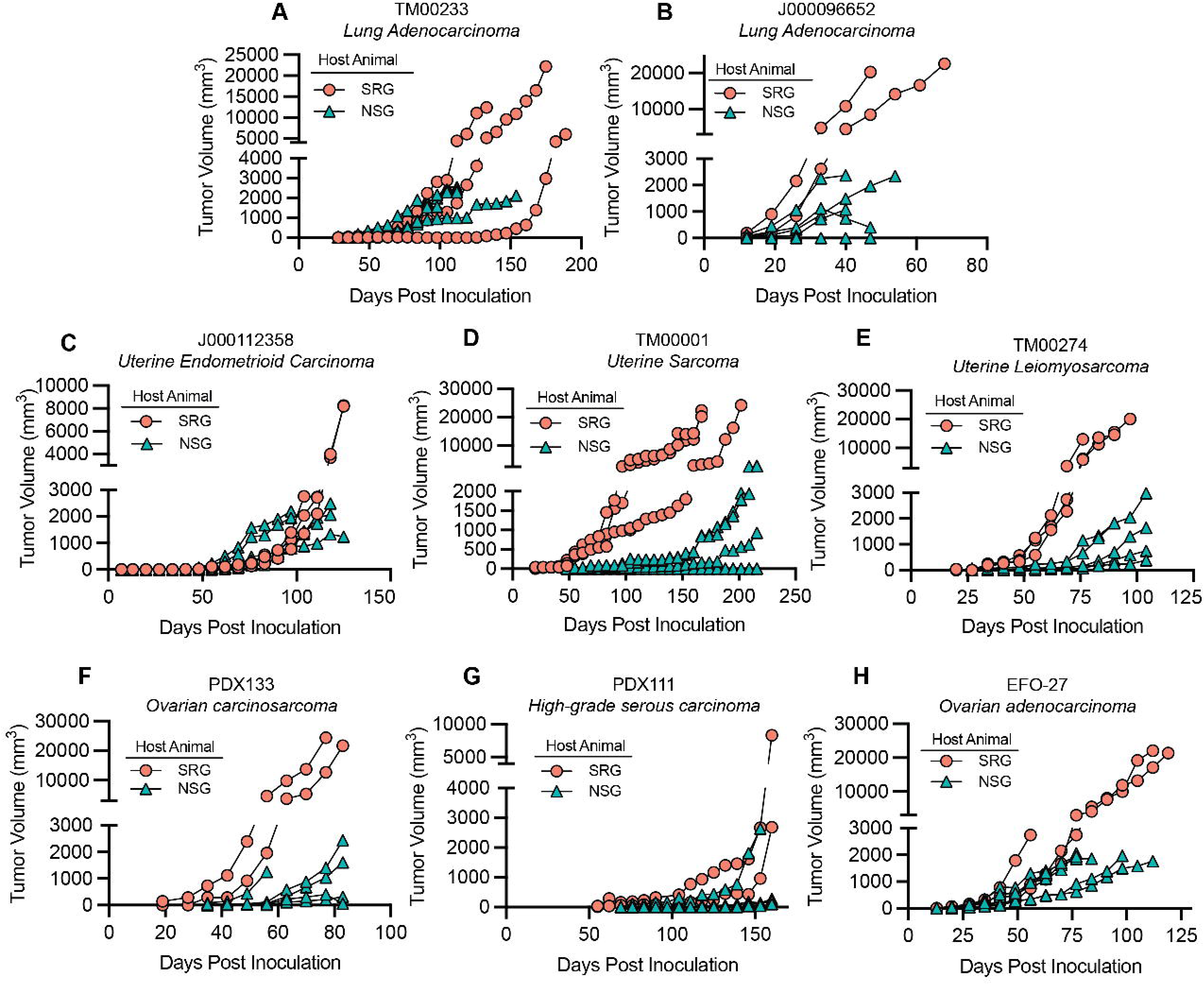
SRG rats support the growth and engraftment of multiple PDX and CDX models over conventional immunodeficient mice. **A,** 8 mm^3^ fragments of lung adenocarcinoma PDX TM00233 were implanted subcutaneously into NSG mice (n=4) or SRG rats (n=3). **B,** 8 mm^3^ fragments of lung adenocarcinoma PDX J000096652 were implanted subcutaneously into NSG mice (n=5) or SRG rats (n=2). **C,** 8 mm^3^ fragments of uterine carcinoma PDX J000112358 were implanted subcutaneously into NSG mice (n=5) or SRG rats (n=3). **D,** 8 mm^3^ fragments of uterine sarcoma PDX TM0001 were implanted subcutaneously into NSG mice (n=5) or SRG rats (n=3). **E,** 8 mm^3^ fragments of uterine leiomyosarcoma PDX TM00274 were implanted subcutaneously into NSG mice (n=5) or SRG rats (n=3). **F,** 8 mm^3^ fragments of high-grade ovarian carcinosarcoma PDX133 were implanted subcutaneously into NSG mice (n=5) or SRG rats (n=2). **G,** 8 mm^3^ fragments of high-grade ovarian serous carcinoma PDX111 were implanted subcutaneously into NSG mice (n=5) or SRG rats (n=2). **H,** 8 mm^3^ fragments of ovarian adenocarcinoma CDX EFO-27 were implanted subcutaneously into NSG mice (n=5) or SRG rats (n=3). For all graphs (**A-H**), Tumor volumes were measured by caliper measurement, and individual animals are plotted in the graph.

**Table 1:**
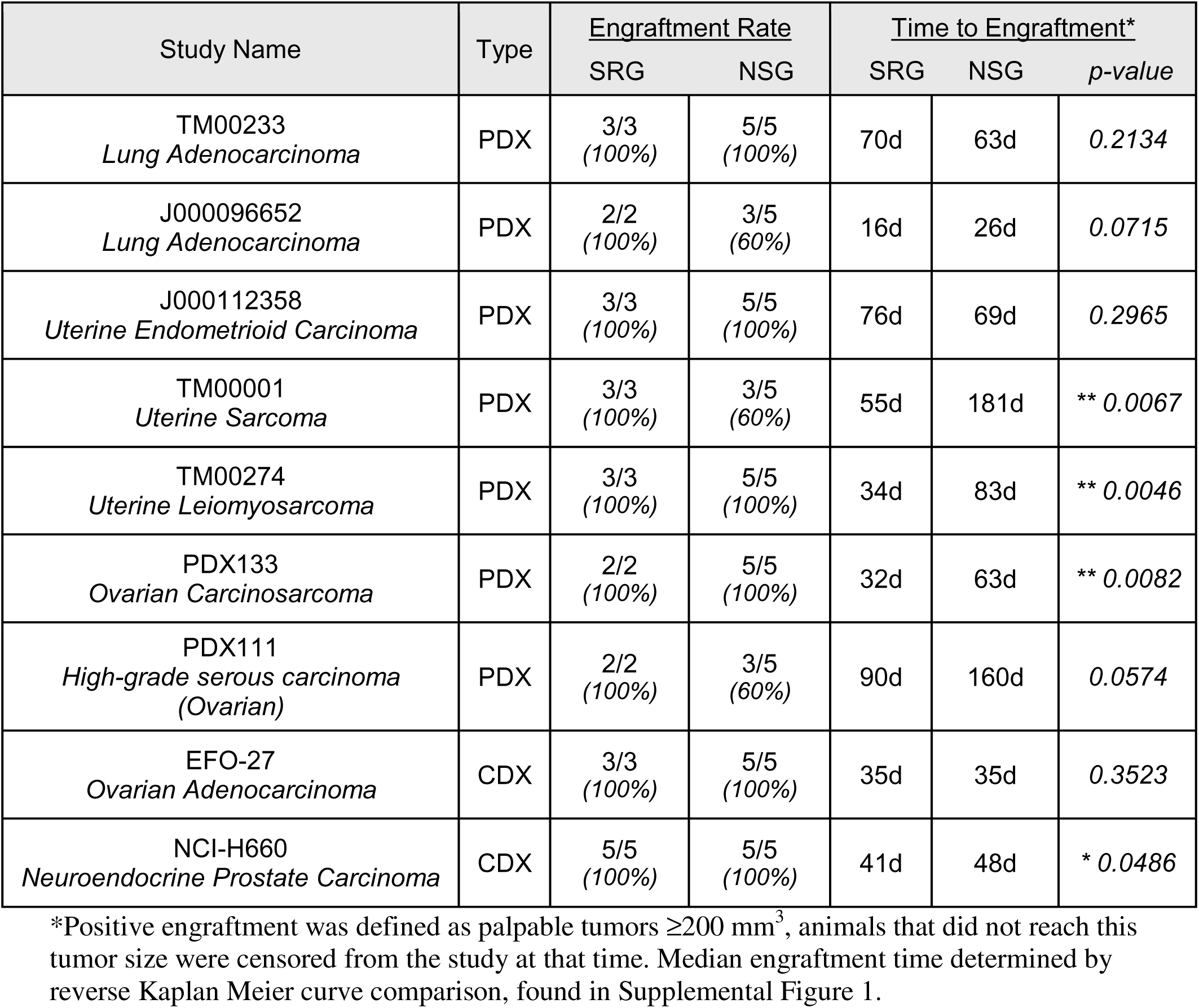
Summary of engraftment rates and time to tumor growth in all PDX and CDX xenografts.

Taken together, all eight models had a 100% engraftment rate in the SRG rats, compared to only four models with a 100% engraftment rate in NSG mice. Additionally, the time to engraftment was either similar between the two species or occurred significantly earlier in the SRG rats. Some models showed a reduction in the time to tumor formation by 50 days or more (**Table 1**). These data are supportive of previous findings that have shown that the SRG rat supports the engraftment and growth of multiple models of human cancer.

### Lung adenocarcinoma PDX models grown in the SRG rat display a microscopic morphology that more closely represents the originating patient tumor

H&E images from the primary patient samples from the lung adenocarcinoma PDX models are available from Jax Lab (**Figure 2A and C**). A comparison of H&E images of primary patient samples and TM00233 and J000096652 grown in NSG mice and SRG rats was performed and showed that in both tumor types, neoplastic cell morphology was comparable, while fibrovascular stroma was denser and more extensively deposited in the SRG rat than the NSG mouse (**Figure 2B and D**). This finding suggests that the TME supported by the SRG rats is morphologically more consistent with the primary patient lung adenocarcinoma samples than the NSG mice.

**Figure 2:**
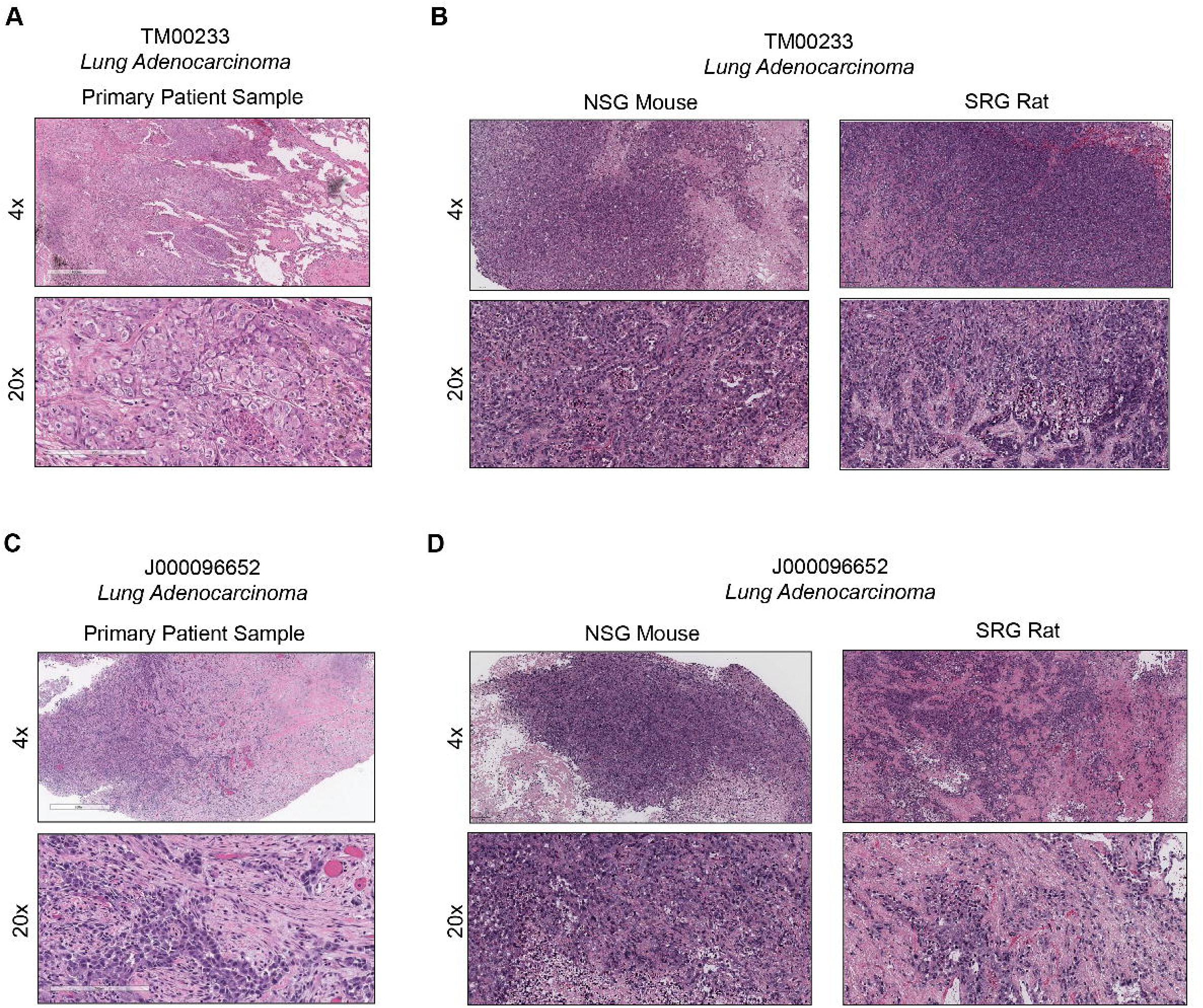
Lung adenocarcinoma PDX models grown in the SRG rat display a microenvironment that more closely represents the originating patient tumor. **A**, H&E images from the primary patient sample where the TM00233 PDX was derived, 4X magnification (top), 20X magnification (bottom). Scale bars indicate 200 μm (20x) and 500 μm (4x). Images from The Jackson Laboratory’s Mouse Tumor Biology Database. **B**, Representative images from TM000233 tumors grown in the NSG mouse (left) or SRG rat (right). 4X magnification (top), 20X magnification (bottom). Scale bars indicate 50 μm (20x) and 250 μm (4x). **C**, H&E images from the primary patient sample where the J000096652 PDX was derived, 4X magnification (top), 20X magnification (bottom). Scale bars indicate 200 μm (20x) and 600 μm (4x). Images from The Jackson Laboratory’s Mouse Tumor Biology Database. **D**, Representative images from J000096652 tumors grown in the NSG mouse (left) or SRG rat (right). 4X magnification (top), 20X magnification (bottom). Scale bars indicate 50 μm (20x) and 250 μm (4x).

### SRG rats support the growth and engraftment of the castrate-resistant prostate cancer CDX model over conventional immunodeficient mice

PDX and CDX models of prostate cancer remain limited and difficult to establish using conventional immunodeficient animals (22). We chose the castrate-resistant NCI-H660 cell line for head-to-head comparison of the growth rate in SRG rats, NSG mice, and NOD-SCID mice recipient animals. Two million NCI-H660 cells were injected into the right flank of 6–8-week-old non-castrated animals in 50% Matrigel. Animals were monitored weekly until tumors were palpable, and then tumors were measured twice weekly by caliper measurement (**Figure 3A**). As with the previous models, cells were prepared and injected simultaneously for all host animals. Interestingly, the NCI-H660 cells had no engraftment or tumor growth in the NOD-SCID mice. The engraftment rates of the NCI-H660 cells were equivalent in the NSG mice and SRG rats at 100% for both species (**Table 1**). However, while the tumor growth plateaued for the NSG mice before humane endpoints, we observed more robust and continuous tumor growth of the NCI-H660 cells in the SRG rat (**Figure 3A**). Additionally, the time to tumor formation was significantly shorter for tumors grown in the SRG rat (**Supplemental Figure 2**).

**Figure 3:**
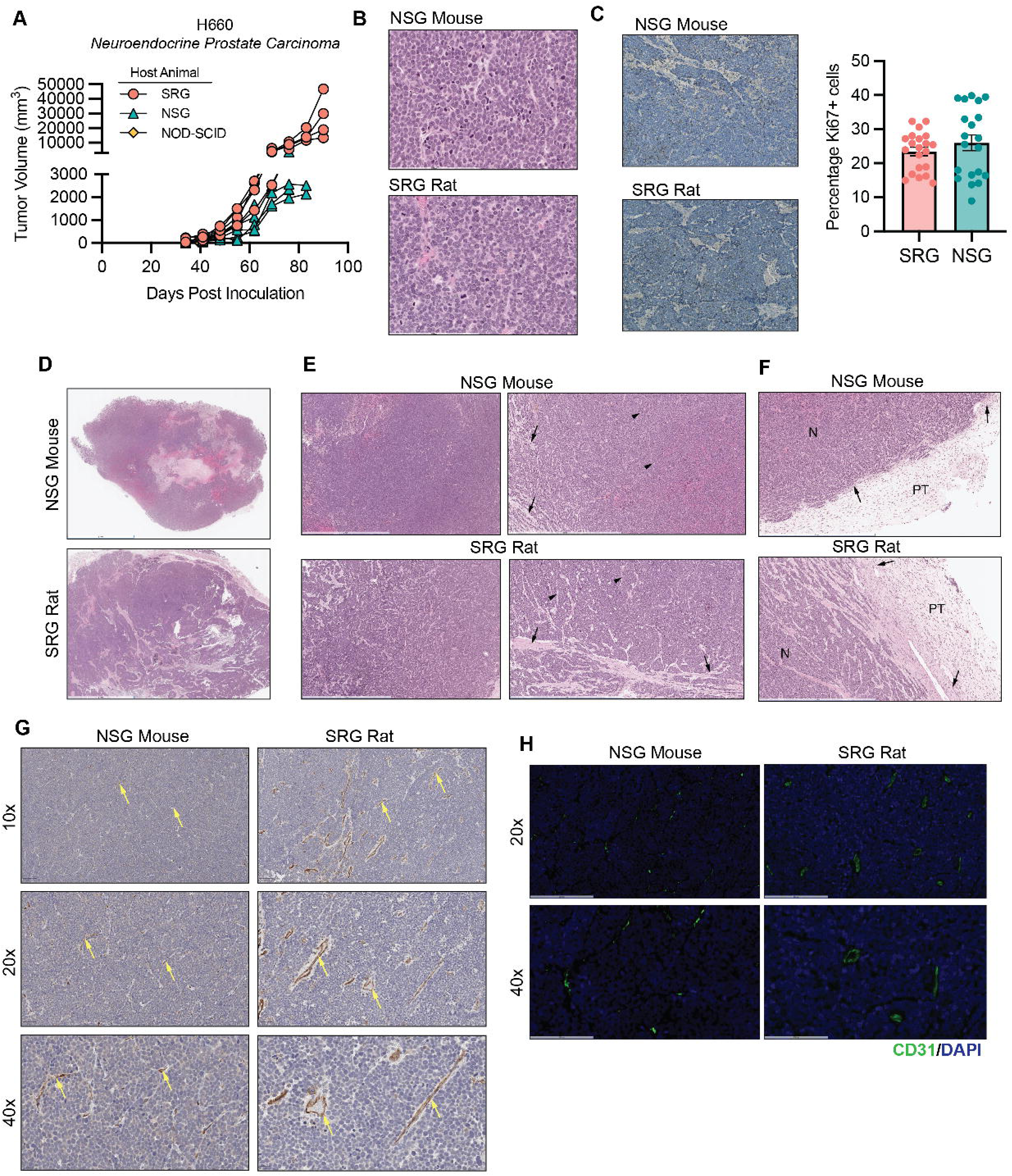
SRG rats support the growth and engraftment of the castrate-resistant prostate cancer CDX model over conventional immunodeficient mice. **A,** NCI-H660 cells were injected subcutaneously into host animals in 50% Matrigel, and growth was tracked by caliper measurement. Each animal was plotted individually. **B,** Tumors from the SRG rat and NSG mouse were fixed, paraffin-embedded, and H&E stained. A representative 400x image is shown, scale bar indicates 200 μm. Pathology determined the morphology and mitotic count of the NCI-H660 cells to be equivalent between the species. Quantification can be found in Supplemental Figure 3. **C**, Representative images (10x images, scale bar indicates 100 μm) and quantification of the Ki67 staining by IHC of H660 tumors; n=4 H660 tumors from each host species were used for the analysis. Error bars ± SD. Larger panel of representative images can be found in Supplemental Figure 4. **D**, Representative H&E image of a 1x cross-section of NCI-H660 tumor grown in the NSG mouse (top) or SRG rat (bottom). Necrosis (black arrows) and viable islands surrounding vascular profiles (black arrowheads). Scale bar indicates 5 mm. **E**, Representative H&E images of tumor stroma at 40x (left) and 100x (right). 100x images show fibrovascular stroma (black arrowheads) and dense collagen/ECM tracts (black arrows). Scale bar indicates 2 mm (40x), 1 mm (100x). **F**, Representative H&E images of capsule formation. Collagen forming a capsule (black arrows), surrounding the neoplasm (N), and peritumoral tissue (PT) indicated. **G**, Chromogenic CD31 staining of immunoreactive blood vessels (DAB chromogen; yellow arrows), tend to be smaller and more numerous in mouse tumor tissue, with larger blood vessels supported by more abundant stroma in rat tumor tissue. Scale bar indicates 100 μm (10x), 50 μm (20x), 20 μm (40x). **H**, Fluorescent CD31 immunoreactive blood vessels (green channel) tend to be smaller and more numerous in mouse tumor tissue, with slightly sparser and larger blood vessels in rat tumor tissue. Scale bars indicate 200 μm (20x), 100 μm (40x). Quantification of G can be found in Supplemental Figure 6.

To explore why the SRG rat supported robust tumor growth of the NCI-H660 CDX model, we performed histology and immunohistochemistry on the four tumors grown in each host species. First, tumor sections were H&E stained, and histopathology interpretation was performed. Interestingly, the NCI-H660 neoplastic cells displayed similar morphologic characteristics and mitotic counts regardless of the host species (**Figure 3B, Supplmental Figure 3**). This was confirmed by IHC using Ki67 staining, which also showed no differences in the percentage of Ki67 positive cells between the host species (**Figure 3C, Supplemental Figure 4**). However, the tumors grown in NSG mice showed more extensive central necrosis that contained hemorrhage, while the tumors grown in the SRG rat showed smaller regions of necrosis throughout the tumor, with viable islands surrounding vascular profiles (**Figure 3D, Supplemental Figure 5**). Additionally, the tumors grown in the SRG rat showed more extensive fibrovascular stroma throughout the tumor (**Figure 3E**, bottom left), as well as thicker, mature tracts of fibrous tissue in some regions (**Figure 3E**, bottom right). In xenograft experiments, a layer of host fibrous tissue can encapsulate the implanted neoplastic cells (23). In tumors from NCI-H660 cells implanted into NSG mice, we observed minimal capsule formation and the surrounding peritumoral tissue lacked significant infiltrating cells (**Figure 3F, top**). Conversely, in the tumors from cells implanted into SRG rats, the capsule was well developed, with matrix and stroma extending into the neoplasm and with more infiltrating cells in the surrounding peritumoral tissue (**Figure 3F, bottom**). Given the greater abundance of fibrovascular stroma in the tumors implanted in SRG rats, we performed CD31 staining by IHC and IF to assess the presence of vascular endothelial cells. Although CD31-positive vascular density was similar in tumors from SRG rats and NSG mice (**Supplemental Figure 6A and B**), the morphology of blood vessels in NSG mice appeared smaller versus the tumors from SRG rats, which showed larger blood vessels supported by thicker stroma; a more mature vascular phenotype (**Figure 3G and H**). This was consistent with a higher microvessel density in tumors from NSG mice compared to the SRG rats (**Supplemental Figure 6C**).

Combined, these data indicate that the TME, specifically the increase in the supporting stroma and vascular maturity in the SRG rat, may contribute to enhanced tumor growth.

### Spatial transcriptomics reveals differences in the global spatial architecture of NCI-H660 tumors grown in SRG vs NSG animals and an upregulation of the protumor genes TCIM and CXCL2

To explore differences in the tumor and host cell interactions and gene expression changes, we performed single-cell spatial transcriptomics using the Xenium platform by 10X Genomics. Briefly, a representative NCI-H660 tumor grown in the SRG rat and NSG mouse was selected for Xenium analysis, and a pre-designed panel of 377 cancer-related genes was analyzed. This was a human-specific gene panel to avoid background from gene expression changes between the host species. Both tumor samples showed an equivalent number of decoded transcripts per 100 μm^2^, however, due to the larger size of the NCI-H660 tumor in the SRG rat, twice as many cells and high-quality decoded transcripts were detected compared to the NSG mouse (**Supplemental Figure 7**). To exclude rodent host cells, k-means clustering was used to distinguish between cells with low transcript counts in the host compared to the tumor tissue, followed by validation using genes highly expressed in the tumors grown in both species (ex, *EPCAM*) and more highly represented in the tumors grown in the SRG (ex, *CXCL2*) (**Figure 4A and B**). Consistent with the H&E staining of the larger cohort, the SRG rat sample included a larger proportion of host stroma cells within the assayed tumor compared to the NSG mouse sample (**Figure 4C**). The remaining human tumor cells for each sample were then merged and clustered to identify 14 clusters within the human tumor population (**Supplemental Figure 8, Supplemental Table S1**). These clusters were overlaid onto the Xenium slides, revealing distinct spatial patterns of gene expression between the H660 tumors dependent on the host animal (**Figure 4D and E**). These clusters were further refined, as some clusters contained few cells. For example, Cluster 13 was almost exclusively found in the H660 cells grown in the NSG mouse, while Cluster 6 was unique to the cells grown in the SRG rat (**Supplemental Table S1**). Differential expression (DE) comparisons were made between all shared clusters between the SRG rat and NSG mouse samples and then between unique and shared clusters within each sample. The number of DE genes in between and within slide comparisons was visualized using an upset plot (**Supplemental Figure 9**), showing that the greatest number of significantly differentially expressed genes were in shared clusters, with the top 10 upregulated genes in the H660 cells grown in the SRG rat from the shared clusters shown in **Figure 4F**. This analysis revealed low transcript counts in the host compared to the tumor tissue. Finally, mapping the transcripts of the top five DE genes onto the SRG tumor showed distinct patterning of expression across the tumors (**Supplemental Figure 10**). Expression of *CXCL2* and *TCIM* appeared to have higher expression in tumor regions with morphologically mature and denser stroma than compared to other regions of the tumor. In a blinded fashion, two regions of stroma dense and stroma poor tissue were selected for further analysis within the SRG tumor (**Supplemental Figure 11**). The heatmap of the transcript levels of *TCIM* and *CXCL2* were overlaid onto the DAPI stain to show the abundant expression of these genes surrounding the stoma (**Figure 4G**).

**Figure 4:**
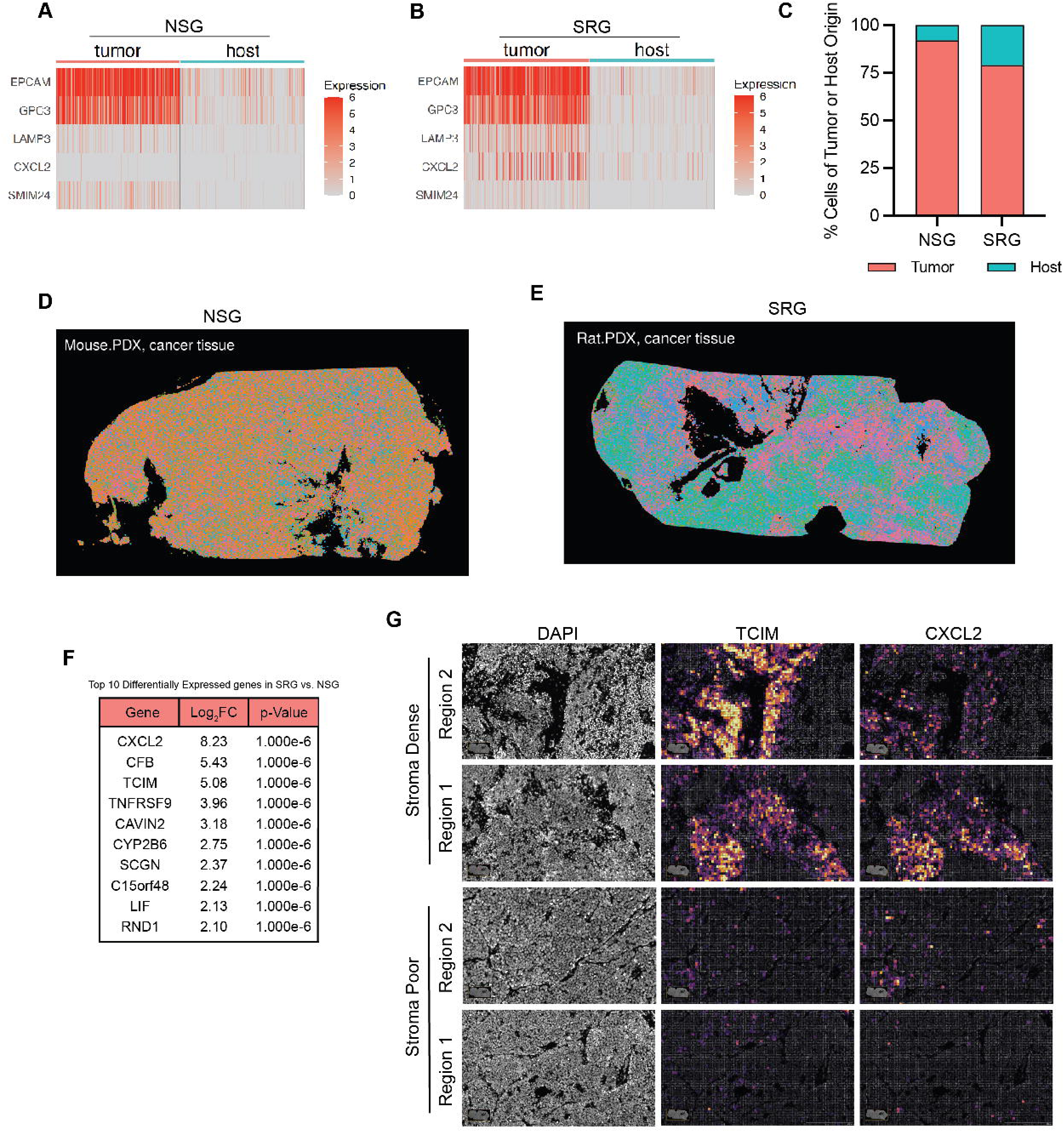
Spatial transcriptomics reveals differences in the global spatial architecture of NCI-H660 tumors grown in SRG vs NSG animals and an upregulation of the protumor genes *TCIM* and *CXCL2*. **A and B,** Expression levels of *EPCAM, GPC3, LAMP3, CXCL2*, and *SIMM24* in host vs tumor cells in the NSG (A) or SRG (B). **C,** Percentage of the total cell counts that were of putative host origin or tumor cell origin. Overlaid images of unintegrated gene expression clustering onto the Xenium slides. **D and E,** Expression levels of *EPCAM, GPC3, LAMP3, CXCL2*, and *SIMM24* in host vs tumor cells in the SRG (E) or NSG (F). **F**, Top ten upregulated DE genes in the H660 tumor cells when grown in the SRG animal. **G,** Spatial mapping of TCIM or CXCL2 gene expression in stroma dense (top) or stroma poor (bottom) tumor regions from the SRG.H&E from these regions is presenting in Supplemental Figure 11. Figure generated in Xenium Explorer 3.2. Scale bars indicate 200 μm.

Taken together, these data were suggestive of host tumor cell interactions leading to gene expression changes in the H660 cells. These gene expression changes were most dramatic in the regions surrounding the stroma, supporting the notion that the microenvironment of the SRG rat may be supporting the growth of the tumor cells.

### Host macrophages are spatially associated with fibrovascular stroma in NCI-H660 tumors

The SRG rat and NSG mouse are similarly highly immunodeficient. The SRG rat has functional deletion of both the *Rag2* and *Il2rg* genes and the NSG mouse has functional deletion of *Prkdc* and *Il2rg*. Both host species are left B and T cell deficient and lack NK cells. However, knockout of these genes will not directly affect macrophages. For this reason, we examined the macrophage populations in the NCI-H660 tumors. First, we performed Iba-1, a macrophage marker, staining by IHC which showed similar macrophage abundance between the NCI-H660 tumors grown in the NSG mice or SRG rats (**Supplemental Figure 12**). However, morphologically, the Iba-1 staining patterns were different between the two species (**Figure 5A**). Specifically, in the NCI-H660 tumor grown in the SRG rat, there were higher levels of Iba-1 staining in areas adjacent to necrosis and areas of abundant stroma. To look more closely at the macrophage populations in the SRG rat tumors, we performed immunofluorescent staining of CD163 and CD68 stroma-poor and stroma-dense regions (Figure 5B). This analysis showed that the CD68-immunolabeled macrophages (green channel) were seen throughout the tumor (T) and were abundant within the adjacent non-tumor stroma (St) (**Figure 5B**). Additionally, the CD163 positive cells, consistent with M2 polarized macrophages, were seen primarily at the border or within the non-tumor stroma (St) and were not as prevalent in stroma-poor regions (**Figure 5B**). Further analysis of the Iba-1-stained tumor sections also showed more robust staining in the stroma-dense regions of the tumor than the stroma-poor tumor regions, which were more prominent in the SRG tumors than the NSG tumors (**Figure 5C**). Taken together, these data indicate that host macrophages are spatially localized to tumor stroma in both species. The NCI-H660 tumors grown in the SRG rat had more abundant stroma, which showed increased expression of tumor-associated macrophages, including M2 polarized macrophages within these regions.

**Figure 5:**
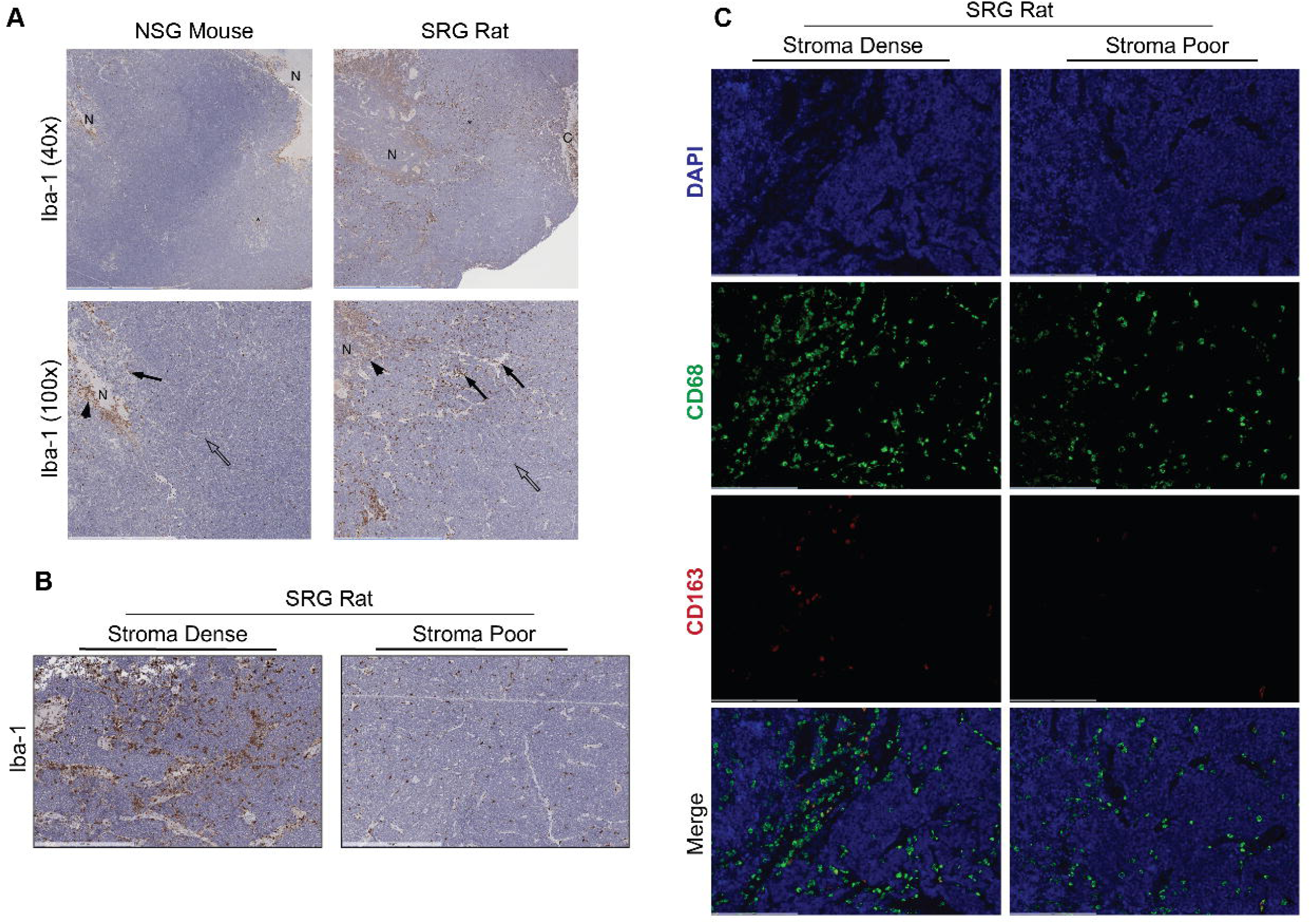
NCI-H660 tumors grown in the SRG rat show activated macrophages in stroma-dense regions. All slides were obtained serially from the section used for Xenium analysis **A**, Iba-1 immunolabeling of macrophages by DAB chromogen in mouse (left) and rat (right) tumors. Macrophages adjacent areas of necrosis (black arrowhead), in areas of abundant stroma (black arrows) and less stroma (open arrows) are shown. Areas of necrosis (N), capsule (C), denser stroma (*) are indicated. Scale bar indicates 2 mm (4x; top row) and 1 mm, 10x; bottom row). Quantification can be found in Supplemental Figure 12. **B**, Iba-1 immunolabeling of macrophages by DAB chromogen in stroma-dense (left) or stroma-poor (right) regions of tumors taken from the SRG rat. Scale bar indicates 500 μm (10x). **C,** CD68/CD163 dual IF for macrophages in stroma-dense and stroma-poor regions of SRG rat. DAPI: top row, CD68 (green channel), CD163 (red channel), Merged image: bottom row; CD68+CD163+ cells are yellow. Scale bar indicates 200 μm (30x).

## Discussion

Low engraftment rates and slow growth limit current PDX and CDX models in immunodeficient mice. Previous work has highlighted the potential benefit of the SRG rat, a new immunodeficient rat strain with the functional deletion of both the Rag2 and Il2rg genes on the Sprague-Dawley background (SRG rat), leading to loss of B, T, and NK cells. This immunodeficient host model has been shown to increase engraftment rates and tumor growth, however, the mechanisms underlying this observation have not been previously explored.

Here, we directly compared the xenotransplantation of multiple PDX and CDX models into the SRG rat and the commonly used immunodeficient mouse strain NSG. We chose NSG mice, as they are a highly immunocompromised strain of mice available and are considered one of the most sensitive hosts for cancer stem cells when compared to NOD-SCID or nude immunodeficient animals (ref). Interestingly, while the SRG rat is not more immunocompromised than the NSG mouse, the growth of the tested PDX and CDX models was better supported in the SRG rat. While our analysis of the NCI-H660 tumors showed no differences in the percent of proliferating cells determined by Ki67 or mitotic count at study endpoint, it is important to note that this does not exclude the potential differences earlier in the study, nor does it exclude potential differences in cell cycle progression rates. Given the size differences in the tumors at the study endpoint and the injection or implantation of the same amount of starting material, it would be important to look at proliferation rates over time in both host animal species.

While NSG mice are considered one of the better immunodeficient models for maintaining human TME (24), a comparison of the microenvironment of the tumors when grown in each host species showed that tumors grown in the SRG rat had more mature stromal compartments and vascular formation, which also mirrored those of the corresponding patient tumors, highlighting that the SRG rat may provide an opportunity to study tumor-stromal interactions. Non-hematopoietic cells in the TME have been shown to produce immunomodulatory factors and cytokines, influencing interactions between the tumor and the microenvironment. Macrophages, including those with M2 polarization, also appeared to be spatially associated with tumor fibrovascular stroma in our models. Interestingly, M2 macrophages have been shown to contribute to the immunosuppressive nature of the TME and facilitate tumor metastasis (25, 26). Additionally, M2 macrophages, like M1 macrophages, can secrete cytokines and chemokines to alter tumor growth, vascularization, and the immune microenvironment (27). Spatial transcriptomics revealed gene expression changes occurring around these stromal compartments. This included the increased expression of *TCIM* and *CXCL2*, both associated with protumor formation and poor prognosis in patients (28–30). Determining the effects of the TME and macrophage populations in regulating these gene expression changes warrants further study.

In summary, the data show that the SRG rat supports the growth of various human cancer types and exhibits improved tumor microenvironment interactions compared to NSG mice. Furthermore, the more developed stromal and vascular compartments of tumors grown in SRG rats were closer to the architecture of the original patient tumors, emphasizing the potential of the SRG rat as a valuable model for human cancer.

## Materials and Methods

### Cell Lines and Cell Culture

NCI-H660 cells were received from Dr. Joshi Alumkal at the University of Michigan and was cultured in RPMI-1640 media supplemented with 0.005 mg/mL insulin, 0.01 mg/mL transferrin, 30nM sodium selenite, 10 nM hydrocortisone, 10 nM beta-estradiol, 2 mM L-glutamine, 5% FBS, and 0.5% penicillin/streptomycin. EFO-27 was purchased from Leibniz Institute DSMZ and was cultured in RPM1 1640 media supplemented with 2 mM L-glutamine, 1X MEM non-essential amino acids, 1 mM sodium pyruvate, 20% FBS, and 0.5% penicillin/streptomycin. All cells were grown in a humidified atmosphere containing 5% CO_2_ at 37°C. All cell lines underwent monthly testing for *Mycoplasma* contamination.

### Patient-derived Xenograft (PDX) Models

TM00233, TM00001, TM00274, J000096652, and J000112358 PDXs were purchased from The Jackson Laboratory. PDX133 and PDX111 were developed and generated by Dr. Analisa DiFeo. Additional clinical information regarding the models can be found in **Supplemental Table S2**. TM00233 was passaged in NOD.Cg-*Prkdc^scid^ Il2rg^tm1Wjl^*/SzJ (NSG mice; Jackson Laboratories) harvested and frozen in 10% DMSO and 90% FBS, then re-implanted study cohort, J000096652, TM00001, TM00274, J000112358, PDX133, and PDX111 were passaged in NSG mice, harvested and implanted fresh into the study cohorts.

### PDX and Cell line derived Xenograft (CDX) studies

All procedures were performed at the University of Michigan and were approved by the University of Michigan Institutional Animal Care and Use Committee. Human-derived tumors were engrafted as a tumor fragment (PDX) or injected in a single cell suspension (CDX) subcutaneously into host SRG RAT^®^ or NSG mice. 6–8-week-old Sprague-Dawley Rag2 null Il2rgamma null ‘SRG’ rats developed by Hera BioLabs, Inc. were purchased from Charles River Laboratories (strain: 707; Sprague Dawley-*Rag2^em2hera^Il2rg^em1hera^/HblCrl*) and 6-8 week old NOD-*scid* IL2Rgamma^null^ (NOD.Cg-*Prkdc^scid^ Il2rg^tm1Wjl^*/SzJ or NSG) mice were purchased from The Jackson Laboratory (strain: 005557). <u>PDX Engraftment</u>: 8 mm^3^ Fresh or frozen tumor fragments were prepared using sterile technique. Tumor preparation and implantation were done at the same time, and the same size of fragments was used for both Mice and Rats. The tumor fragments were engrafted bilaterally in both flanks or unilaterally into the right flank using a trocar needle. <u>CDX Injection</u>: Cells were trypsinized, counted, and suspended in 50% phosphate-buffered saline and 50% Matrigel (Corning Reduced Growth Factor 354230). 2×10^6^ NCI-H660 cells per mouse or rat were injected in 100 μL, and 1×10^7^ EFO-27 cells per mouse or rat were injected in 200 μL for the published studies. Cell injections were performed using a 26g needle. Tumor volume was calculated as (L x W^2^/2), where length and width were measured with digital calipers. Animals were allowed to reach the endpoint of 4 cm (rats) or 2 cm (mice) in any direction. Upon reaching the humane endpoint, the animals were euthanized, and tumors were snap-frozen or formalin-fixed for additional analysis.

### Xenium

FFPE tissue blocks were sent to the Advanced Genomics Core at the University of Michigan for processing using 10x Genomics demonstrated protocols (CG000578, CG000580, CG000582, CG000613). Tissue sections (5 μm) were collected onto Xenium slides where sections were deparaffinized and decrosslinked. RNA within the tissue was labeled using circularizable DNA probes targeting 377 genes included in the predesigned Human Multi-Tissue and Cancer panel (10x genomics, 1000626). Following ligation, the DNA probes were enzymatically amplified followed by autofluorescence quenching and nuclei staining with DAPI. Xenium slides were loaded into the Xenium Analyzer instrument for imaging and analysis, where fluorescently labeled oligos bound the amplified DNA probes, and samples underwent successive rounds of fluorescent probe hybridization, imaging, and removal to generate optical signatures which were converted into a gene identity. Following the Xenium run, slides were H&E stained, and high-resolution images captured for inclusion in the final data set. Acquired data was transferred off the Xenium instrument and data visualized using the Xenium Explorer (10x desktop software) and third-party analysis tools.

### Xenium data analysis

Cell segmentation, gene transcript by cell, and transcript by tissue location data matrices were generated by the Xenium Onboard Analysis pipeline. All downstream analysis and graphics were generated in R (v 4.4.0) (31) and performed primarily using the Seurat package (v 5.0.1) (32). For each slide, automated clustering results for k-means = 2 was used to distinguish between engrafted human tumor cells and host cells. After validation, tumor cells were further filtered to 20-800 counts/cell and a minimum of 5 features/cell.

Expression counts were then normalized using the SCTransform method (33). Normalized data were then merged and standard clustering was performed with 11 PCs and 0.4 resolution was used to generate distinct populations and visualized by uniform manifold approximation and projection (UMAP)(34). To identify comparable tumor populations between slides, clusters were further categorized as “shared”, if at least 3000 cells were represented in that cluster for both slides. MAST (35) was then used to identify differentially expressed (DE) genes, with an adjusted p-value threshold for shared clusters between slides with absolute fold-change > 0.25 and adjusted p-value < 0.01. Functional analysis, including candidate pathways activated or inhibited in comparison(s) and GO-term enrichments, was performed using iPathway Guide (Advaita) (36).

### Immunohistochemistry

CD31 and Iba1 immunohistochemistry (IHC) was performed on one mouse and rat slide each sectioned in a serial manner (4μm) with the slide collected for Xenium analysis, by DTR Labs; all staining was performed on the Leica Bond RX automated stainer (Leica Biosystems, Inc. [Buffalo Grove, IL, USA]).

**CD31**: For the CD31 IHC, slides were heated for 20 minutes at 100°C in pH9.0 EDTA buffer for antigen retrieval. Primary antibody incubation (1:500, rabbit mAb [EPR17259], Abcam #182981) was performed at room temperature for 30 minutes. An HRP-conjugated anti-goat secondary antibody polymer was used for primary binding detection with visualization via diaminobenzidine (DAB) application. Hematoxylin counterstain was used to visualize nuclei.

**Iba-1:** For the IHC targeted to the ionized calcium-binding adaptor molecule 1 (Iba-1) antigen, slides were heated for 20 minutes at 100°C in pH9.0 EDTA buffer for antigen retrieval. Primary antibody incubation (1:5000, rabbit mAb [EPR16589], Abcam #178847) was performed at room temperature for 30 minutes. An HRP-conjugated anti-goat secondary antibody polymer was used for primary binding detection with visualization via diaminobenzidine (DAB) application. Hematoxylin counterstain was used to visualize nuclei.

**Ki67**: Four rat xenograft blocks and four mouse xenograft blocks were sectioned at 4um and stained by immunohistochemistry (IHC) targeted to the Ki67 proliferation protein marker. First paraffin was dissolved from the slides by dipping slides in 3 xylene baths for 5 minutes each, 2 baths of 100% ethanol for 5 minutes each, 1 bath of 95% ethanol for 5 minutes, 1 bath of 70% ethanol for 5 minutes then deionized water for 5 minutes. Heat-Induced antigen retrieval (HIER) was performed using pH 6 Citrate Buffer for 20 minutes in a microwave at power 7. Slides were cooled, then boundaries were drawn around each tissue section using a hydrophobic Dako pen, 3% H_2_O_2_ was added to the sections for 10 minutes to block any endogenous peroxidase activity, 5% Goat Serum was added for 30 minutes as a protein block and the Ki67 primary antibody (rabbit mAB D3B5 Cell Signaling #9129) was added to each section at a 1:1000 dilution in 5% Goat Serum and left overnight in a humidified chamber at 4 degrees Celcius. The following day, an HRP-conjugated anti-rabbit secondary antibody polymer was used for primary binding detection with visualization via diaminobenzidine (DAB) application from the Abcam Rabbit Specific HRP/DAB Detection IHC Kit (ab64261). Gill’s Formula Hematoxylin counterstain was used to visualize nuclei.

**Quantification of IHC by Image Analysis Methods**: Whole slide images from CD31 and Iba-1 chromogenic IHC slides were imported into Visiopharm for further analysis by DTR Labs. For both IHC stains, an initial deep learning algorithm trained to detect tissue and exclude white space was applied. Tissue/histologic artifacts were excluded, and large areas of necrosis were manually annotated by the analyst with guidance by a board-certified veterinary pathologist to avoid detection of non-specific immunostaining of necrotic debris; all data are reported within areas of viable tumor only.

For CD31 quantification, immunolabeled endothelial cells forming vascular strcutures was identified by another DLA trained to identify and separate each blood vessel into an isolated label for discrete count quantification. Vascular density was calculated as the number of blood vessels detected, divided by the tumor area (cells number/mm^2^).

For Iba-1 quantification, an algorithm was applied using traditional thresholding techniques based on the chromogenic substrate, DAB. Percent Iba-1-immunolabeled areawas calculated by taking the quotient of detected Iba-1+ area over the total tumor area.

For the Ki67 quantification, 14-21 images were taken of each slide and imported into QuPath. DAB positive cells were determined using QuPath’s positive cell detection command and after manually reviewing detection boundaries. The percentage of Ki67 positive cells out of total cells for each image was recorded and the average of those values was reported for each of the animals and graphed in GraphPad Prism software.

### Immunofluorescence

CD31 and CD68/CD163 dual immunofluorescence (IF) was performed on one mouse and rat slide each, sectioned in a serial manner (4μm) with the slide collected for Xenium analysis, by DTR Labs; all staining was performed on the Leica Bond RX automated stainer (Leica Biosystems, Inc. [Buffalo Grove, IL, USA]).

**CD31**: For immunofluorescence (IF) targeted to the CD31 antigen, slides were heated for 20 minutes at 100°C in pH9.0 EDTA buffer for antigen retrieval. Primary antibody incubation (1:250, rabbit mAb [EPR17259], Abcam #182981) was performed at room temperature for 30 minutes. An anti-rabbit secondary antibody conjugated with AlexaFluor-568 was used for primary binding detection. DAPI counterstain was used to visualize nuclei.

**CD68+CD163**: For dual CD68 and CD163 IF, slides were heated for 20 minutes at 100°C in pH9.0 EDTA buffer for antigen retrieval. Primary antibody incubation (CD68: 1:300, mouse mAb [ED1], Novus Biologicals #NB600-985; CD163: 1:500, rabbit mAb [EPR19518], Abcam #ab182422) was performed at room temperature for 30 minutes. An anti-mouse secondary antibody conjugated with AlexaFluor-647 (for CD68) and an anti-rabbit secondary antibody conjugated with AlexaFluor-750 (for CD163) was used for primary binding detection. DAPI counterstain was used to visualize nuclei.

**Scanning**: Chromogenic IHC slides (CD31 and Iba-1) were scanned on the Leica Aperio AT2 (Leica Biosystems Imaging [Vista, CA, USA]), and the fluorescent slides were scanned on the Zeiss Axioscan 7 (Zeiss Microscopy [Jena, Germany]). (NIH S10OD032267 to Denise Ramirez).

### Morphologic Pathology

H&E-stained slides from SRG rat and NSG mouse CDX and PDX tumors were evaluated by an ACVP-board certified veterinary pathologist (JRD), including determination of H660 tumor mitotic counts, in which neoplastic cell mitotic figures were counted in ten, 0.25mm^2^ fields (avoiding areas of necrosis, stroma, and inflammatory cells) and averaged per sample (n=5/species). Original patient tumors images available from The Jackson Laboratory were reviewed for comparison to rat and mouse PDX tumors.

## Author Contributions

CMO and GN completed the initial manuscript. CMO, KPZ, JRD, KB, GW, GH, FKN, MH, DB, TB, and KP completed experiments and data collection. DK performed statistical analysis. CMO, AD, MJS, GN all contributed to the study design. MJS and GN provided funding support. All authors reviewed and revised the manuscript.

## Data Availability

Code for downstream Xenium analysis is available via GitHub (https://github.com/umich-brcf-bioinf-projects/Narla_gnrla_SP1-Xenium)

## Acknowledgments

Research reported in this publication was supported by Hera Biolabs, DTR Labs, and the National Cancer Institutes of Health under Award Number P30CA046592 using the following Cancer Center Shared Resource: Single Cell and Spatial Analysis Shared Resource. We would also like to acknowledge support from the Bioinformatics Core of the University of Michigan Medical School’s Biomedical Research Core Facilities (RRID: SCR_019168).

