## Supplemental Data for "The SRG RAT® supports human cell xenotransplantation through enhanced tumor microenvironment interactions"

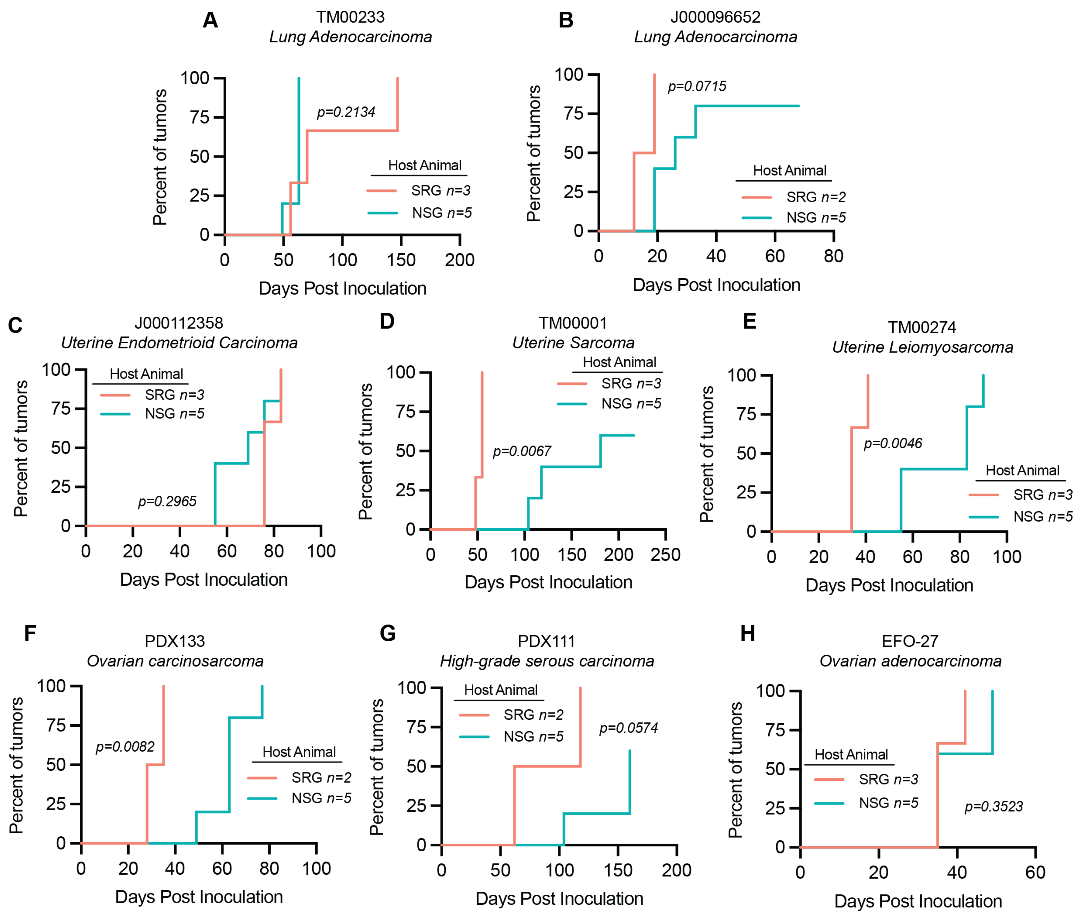


**Supplemental Figure 1: Time to tumor engraftment for PDX and CDX studies.** For all graphs (**A-H**), Tumor volumes were measured by caliper measurement. Positive engraftment was considered when tumors were palpable and reached ≥200 $mm^3$. If tumors were under 200 $mm^3$ at the end of the study, animals were censored at that time. **A,** 8 $mm^3$ fragments of lung adenocarcinoma PDX TM00233 were implanted subcutaneously into NSG mice (n=4) or SRG rats (n=3). **B,** 8 $mm^3$ fragments of lung adenocarcinoma PDX J000096652 were implanted subcutaneously into NSG mice (n=5) or SRG rats (n=2). **C,** 8 $mm^3$ fragments of uterine carcinoma PDX J000112358 were implanted subcutaneously into NSG mice (n=5) or SRG rats (n=3). **D,** 8 $mm^3$ fragments of uterine sarcoma PDX TM0001 were implanted subcutaneously into NSG mice (n=5) or SRG rats (n=3). **E,** 8 $mm^3$ fragments of uterine leiomyosarcoma PDX TM00274 were implanted subcutaneously into NSG mice (n=5) or SRG rats (n=3). **F,** 8 $mm^3$ fragments of high-grade ovarian carcinosarcoma PDX133 were implanted subcutaneously into NSG mice (n=5) or SRG rats (n=2). **G,** 8 $mm^3$ fragments of high-grade ovarian serous carcinoma PDX111 were implanted subcutaneously into NSG mice (n=5) or SRG rats (n=2). **H,** 8 $mm^3$ fragments of ovarian adenocarcinoma CDX EFO-27 were implanted subcutaneously into NSG mice (n=5) or SRG rats (n=3). For all graphs (**A-H**), p-values were determined by Log-rank (Mantel-Cox) test.

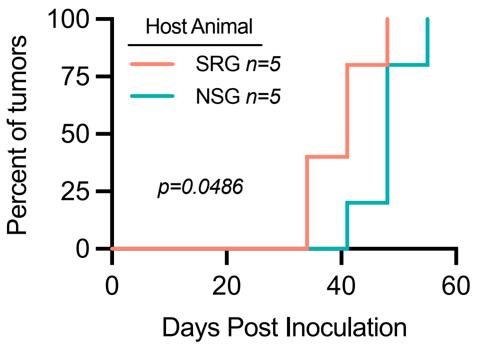


**Supplemental Figure 2: Time to tumor formation for the NCI-H660 xenograft between SRG and NSG animals**. NCI-H660 cells were injected subcutaneously into host animals in 50% Matrigel, and growth was tracked by caliper measurement. Engraftment was considered positive when tumors were palpable and reached ≥200 $mm^3$.  p-values determined by Log-rank (Mantel-Cox) test.

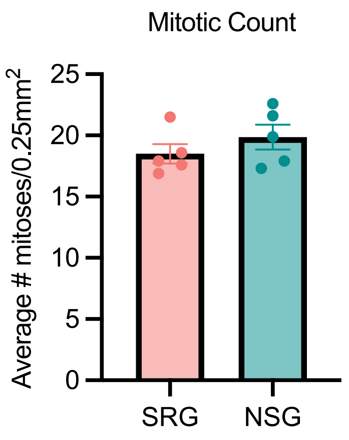


**Supplemental Figure 3: Mitotic Counts from NCI-H660 tumors grown in NSG or SRG animals**.

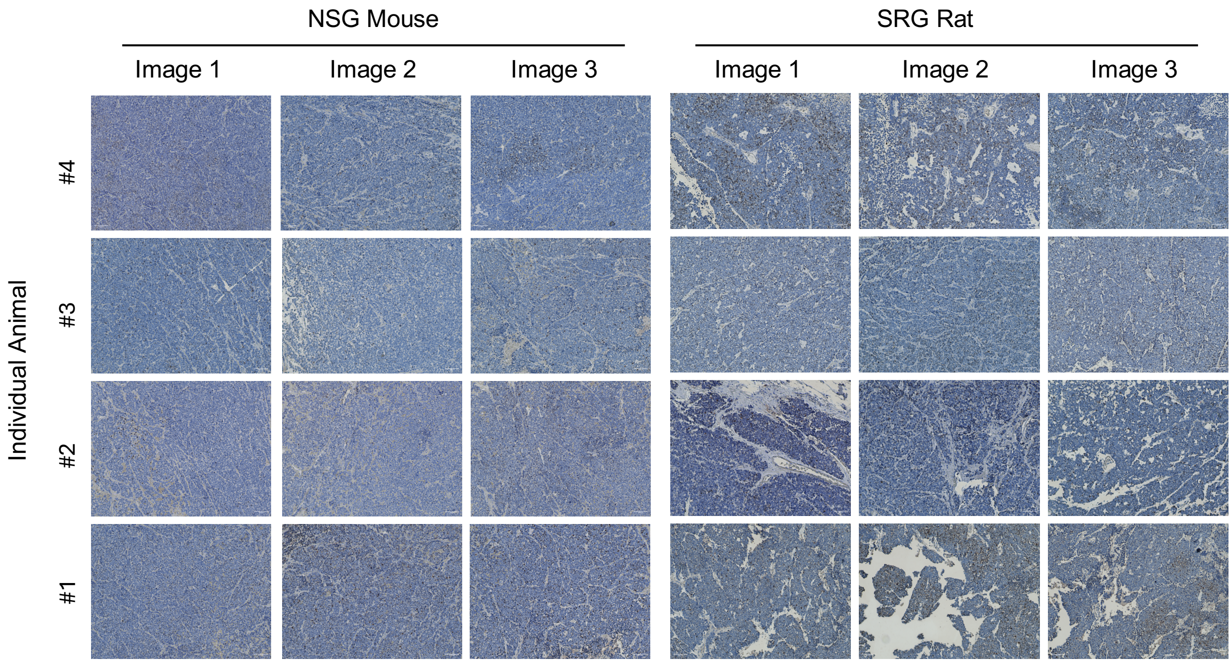


**Supplemental Figure 4: Ki67 Staining from NCI-H660 Animals.** Representative images from Ki67 staining of NCI-H660 tumors grown in NSG mice (left) or SRG rats (right). Staining was performed on 4 individual tumors grown in each host species. Three representative images from each tumor are shown. 10x images, scale bars indicate 100 µm.

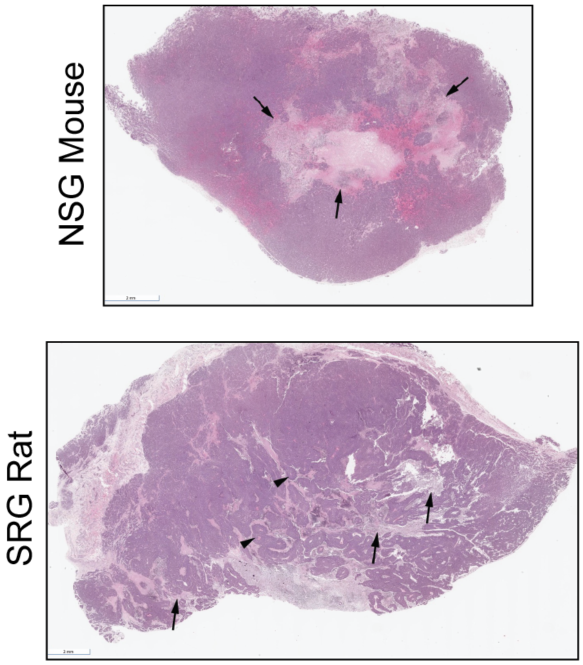


**Supplemental Figure 5: Full Cross-Section of H660 tumors**. H&E image of a ~1x cross-section of NCI-H660 tumor grown in the NSG mouse (top) or SRG rat (bottom). Necrosis (black arrows) and viable islands surrounding vascular profiles (black arrowheads). Scale bar indicates 2 mm.

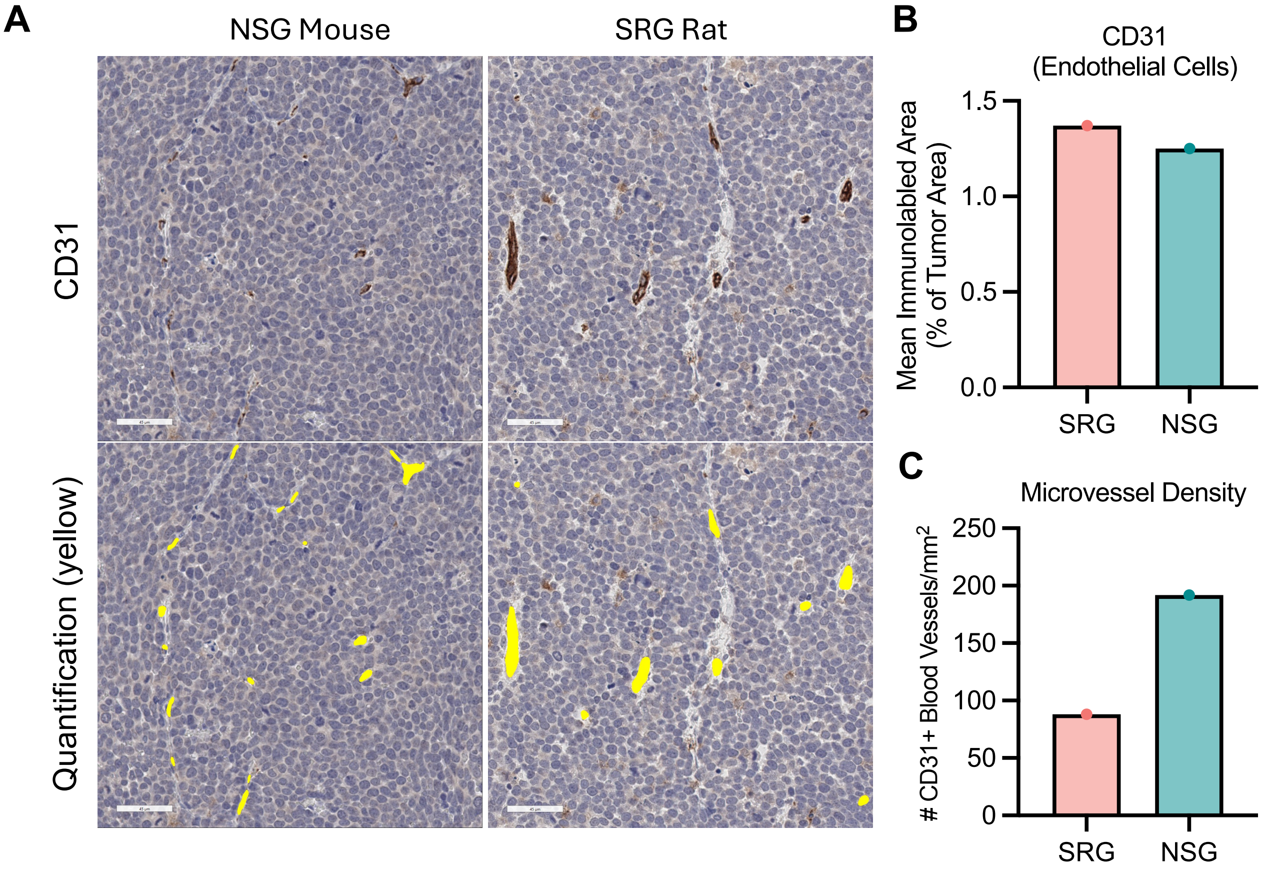


**Supplemental Figure 6: Quantification of CD31**. **A**, CD31 immunoreactive blood vessel (DAB chromogen) in NCI-H660 tumors grown in the NSG mouse, or the SRG rat. Yellow label indicates CD31-immunolabeled blood vessels detected by the algorithm. Scale bar indicates, 45 µm (40X). **B**, Measurement of the overall immunolabeled area is similar between the NSG and SRG animals. The density is higher in mouse compared to rat tissue, while the rat blood vessels tend to be larger. **C**, The vascular density is higher in the mouse compared to rat tissue, while the rat blood vessels tend to be larger.

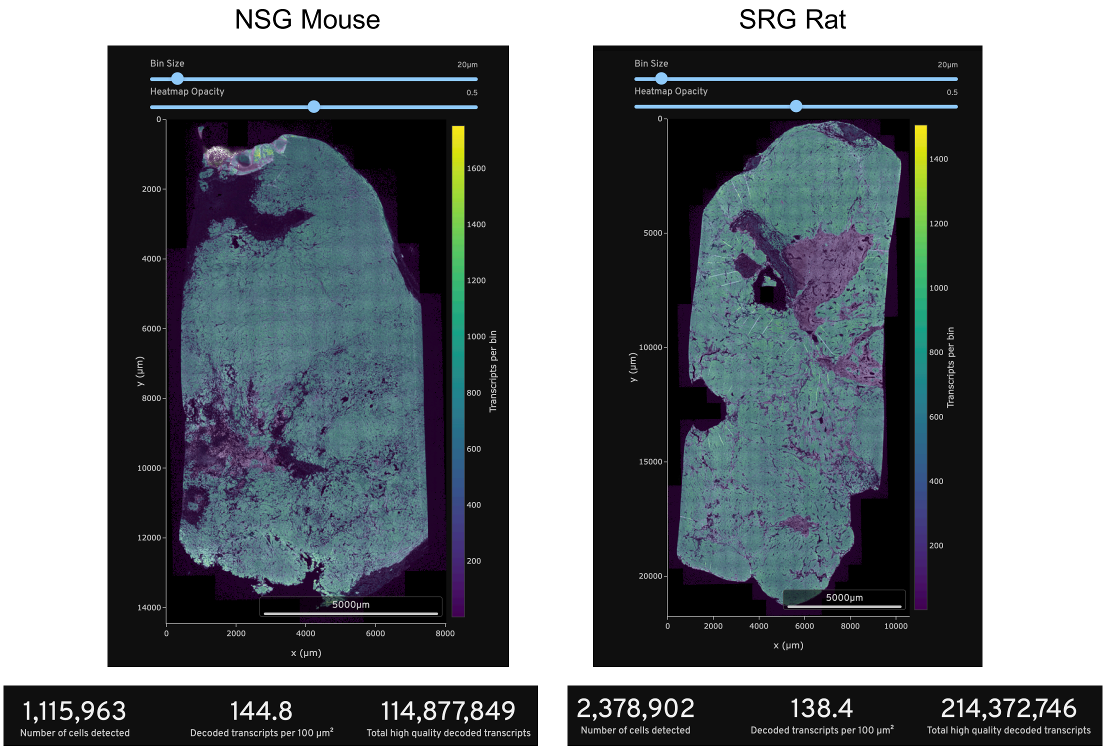


**Supplemental Figure 7: Summary of key metrics from the Xenium analysis**. Images of the cross section of tumor tissue from the NCI-H660 tumor grown in the NSG mouse (left) or SRG rat (right). Below are the number of cells detected, the number of transcripts per 100 μm$^2$, and the total number of transcripts detected in the samples. Scale bars represent 5000 μm.

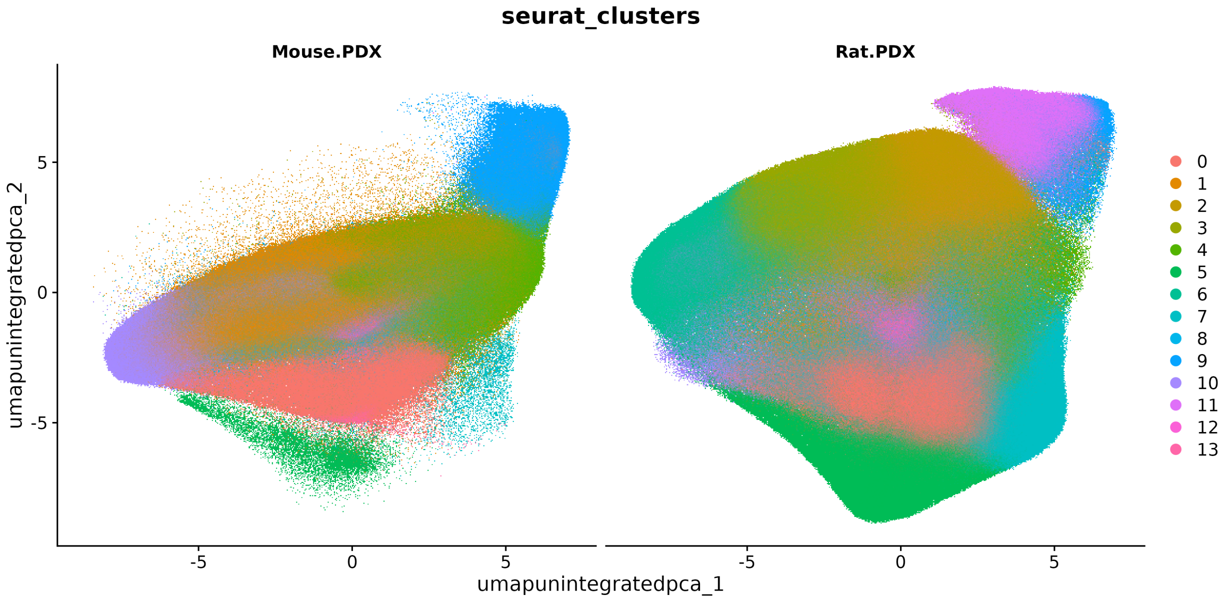


**Supplemental Figure 8: UMAP analysis of unintegrated clusters**. Numerical cell counts for each cluster found in Supplemental Table S1.

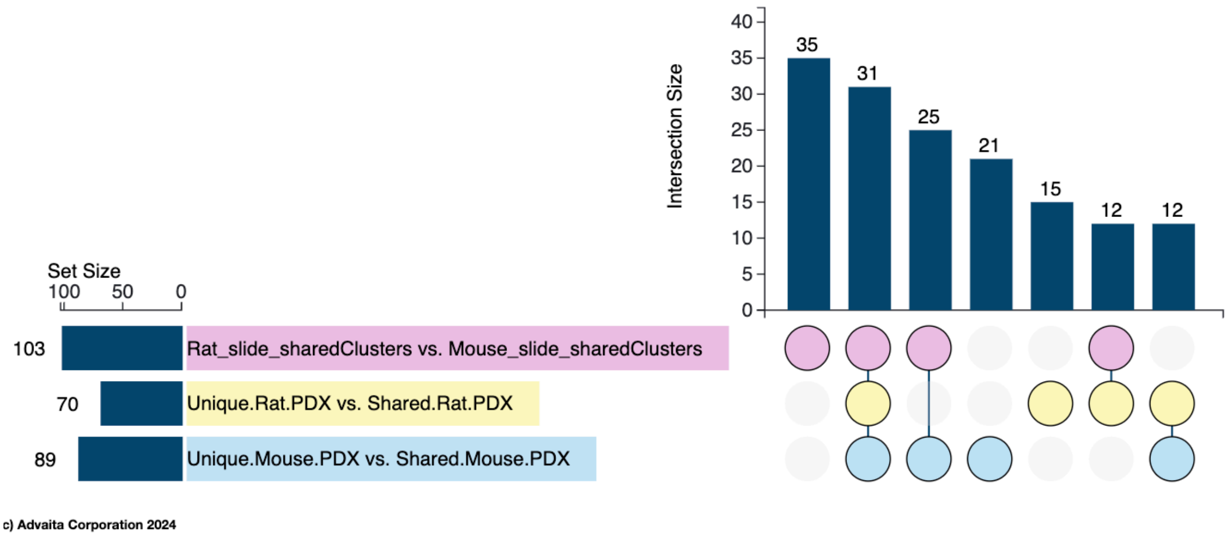


**Supplemental Figure 9**: Upset blot of clusters that were shared or unique to the H660 cells when grown in each host species, related to Supplemental Table S1.

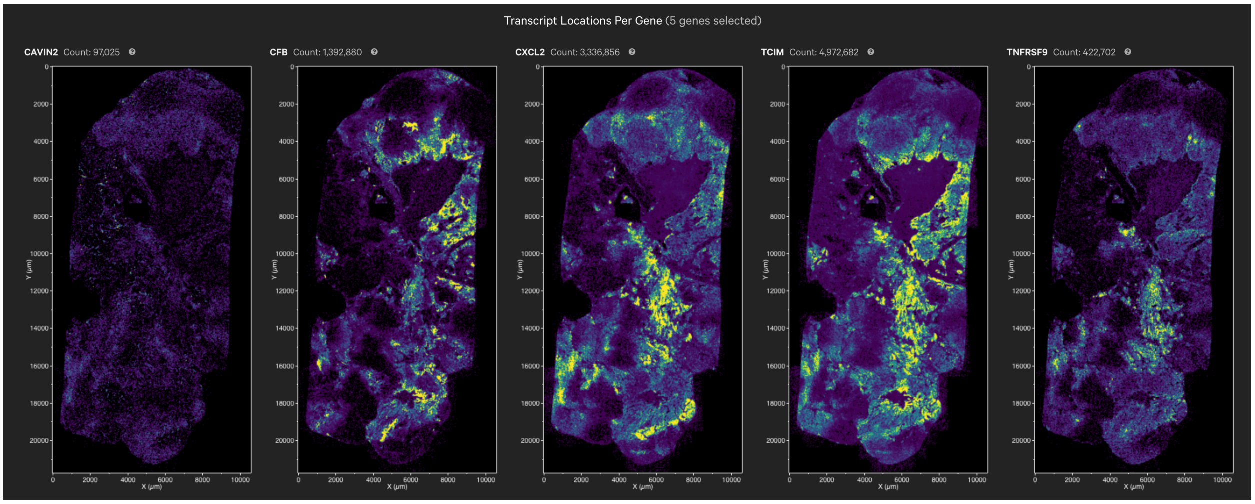


**Supplemental Figure 10: Transcript locations of the genes most upregulated in H660 cells grown in the SRG**. From left to right: *CAVIN2, CFB, CXCL2, TCIM, and TNFRSF9* transcripts were mapped onto the tumor. Figure generated in Xenium Explorer 3.2.

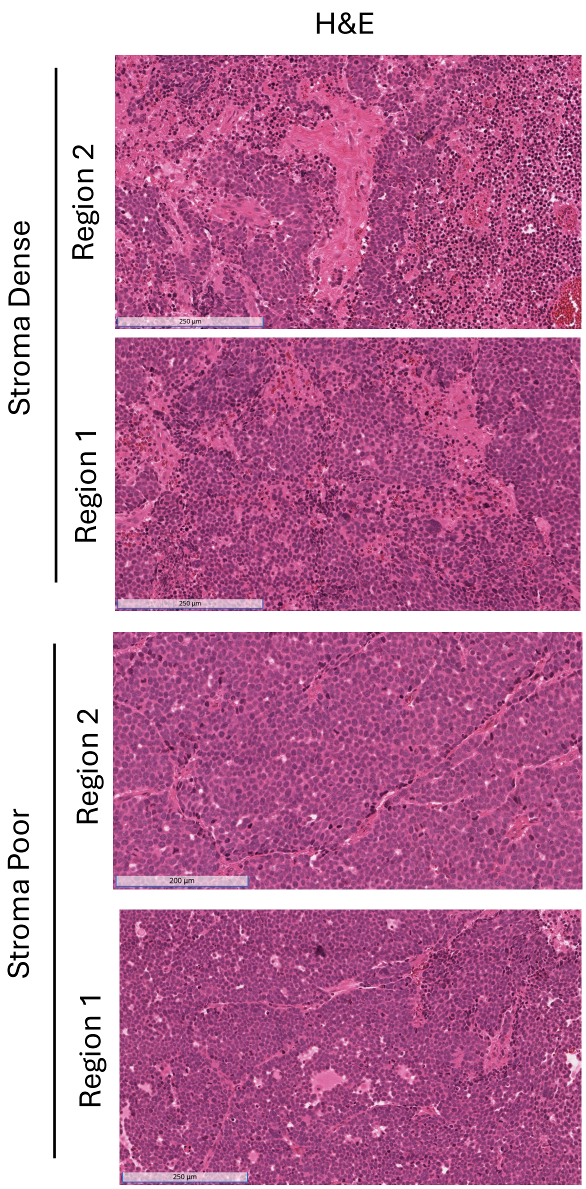


**Supplemental Figure 11. H&E images corresponding to 10x Xenium data.** Scale bars indicate 250 µm.

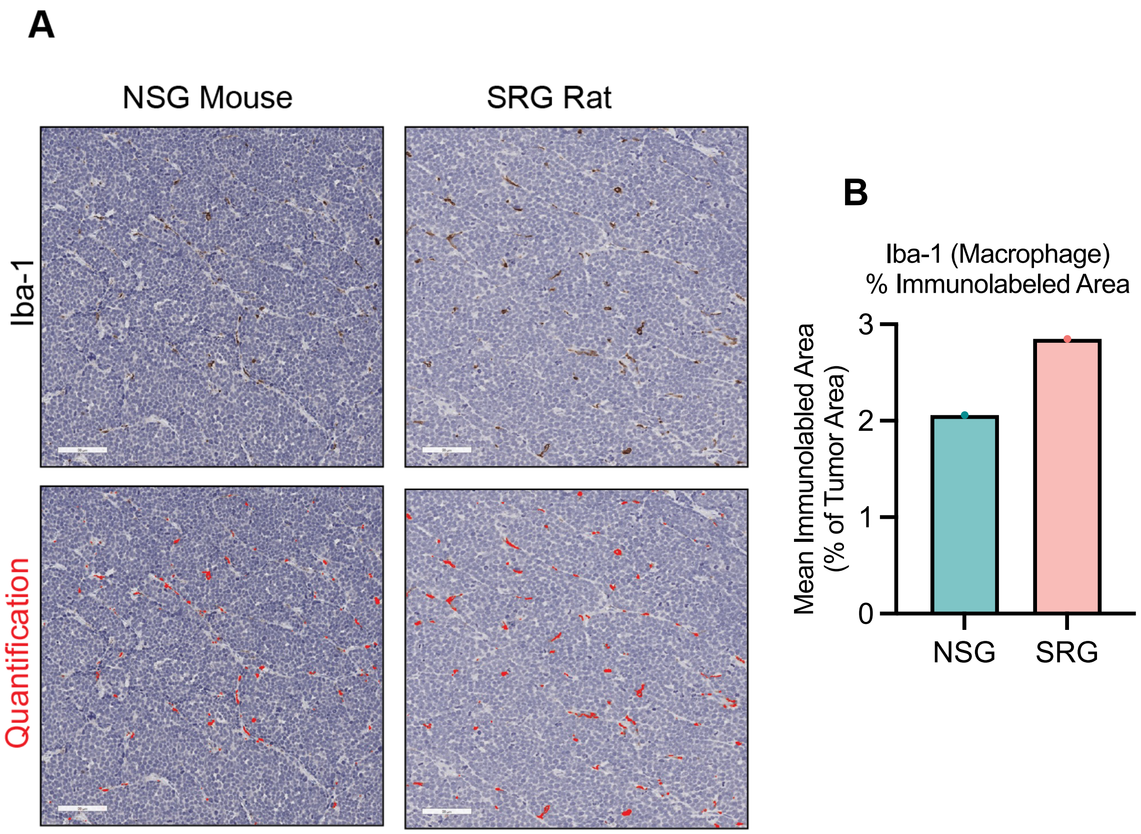


**Supplemental Figure 12. Iba-1 IHC quantification in NSG mouse and SRG rat NCI-H660 tumors.** Slides obtained serially from section used for Xenium analysis **A**, Iba-1 immunolabeling of macrophages by DAB chromogen across all tumor tissue was comparable or slightly greater between the mouse and rat samples. Red label indicates Iba-1-immunolabeled macrophages detected by the algorithm. Scale bar indicates 90 µm (20x). **B**, Quantification of Iba-1-immunolabeled area.

**Supplemental Table S1: Cell Counts per Unintegrated Cluster**

| ClusterID | NSG cell count | SRG rat cell count |
|---|---|---|
| **0** | 113,601 | 288,569 |
| **1** | 357,786 | 43,987 |
| **2** | 1,570 | 314,045 |
| **3** | 388 | 243,720 |
| **4** | 192,646 | 51,066 |
| **5** | 12,971 | 197,243 |
| **6** | 261 | 195,662 |
| **7** | 6,853 | 184,061 |
| **8** | 67,964 | 89,745 |
| **9** | 116,053 | 23,213 |
| **10** | 111,504 | 22,252 |
| **11** | 126 | 123,892 |
| **12** | 18,735 | 28,959 |
| **13** | 2,351 | 25 |

**Supplemental Table S2: Clinical Information from PDX Models**

| Model | Cancer Type | Sample Site | Tumor Type | Stage/Grade | Source |
|---|---|---|---|---|---|
| TM00233 | Lung Adenocarcinoma | Lung | Primary | AJCC IIIA / Grade 3 | The Jackson Laboratory |
| J000096652 | Lung Adenocarcinoma | Lung | Primary | AJCC IIIB / Grade 3 | The Jackson Laboratory |
| J000112358 | Uterine endometrioid carcinoma | Uterus | Primary | AJCC II / Grade 3 | The Jackson Laboratory |
| TM00001 | Uterine Sarcoma | Abdominal Wall | Metastatic | AJCC IV/ Grade 4 | The Jackson Laboratory |
| TM00274 | Uterine Leiomyosarcoma | Lung | Metastatic | AJCC IV | The Jackson Laboratory |
| PDX111 | Ovarian high grade serous carcinoma | Ovary | Primary | AJCC IIIC | Dr. Analisa DiFeo |
| PDX133 | Ovarian high grade carcinosarcoma | Ovary | Primary | AJCC IIIB | Dr. Analisa DiFeo |
